# Near Infrared Reflectance Spectroscopy Phenomic and Genomic Prediction of Maize Agronomic and Composition Traits Across Environments

**DOI:** 10.1101/2023.08.21.554202

**Authors:** Aaron J. DeSalvio, Alper Adak, Seth C. Murray, Diego Jarquín, Noah D. Winans, Daniel Crozier, William Rooney

**Affiliations:** Interdisciplinary Graduate Program in Genetics and Genomics (Department of Biochemistry and Biophysics), Texas A&M University, College Station, TX 77843-2128, USA; Department of Soil and Crop Sciences, Texas A&M University, College Station, TX, 77843-2474, USA; Department of Agronomy, University of Florida, Gainesville, FL, 32611, USA

**Keywords:** Genomic prediction, phenomic prediction, NIRS, yield prediction, kernel composition traits

## Abstract

For nearly two decades, genomic selection has supported efforts to increase genetic gains in plant and animal improvement programs. However, novel phenomic strategies helping to predict complex traits in maize have proven beneficial when integrated into across– and within-environment genomic prediction models. One phenomic data modality is near infrared spectroscopy (NIRS), which records reflectance values of biological samples (e.g., maize kernels) based on chemical composition. Predictions of seven maize agronomic traits and three kernel composition traits across two years (2011-2012) and two management conditions (water stressed and well-watered) were conducted using combinations of NIRS and genomic data within four different cross-validation prediction scenarios. In aggregate, models incorporating NIRS data alongside genomic data improved predictive ability over models using only genomic data in 5 of 28 trait/cross-validation scenarios for across-environment prediction and 15 of 28 trait/environment scenarios for within-environment prediction, while the model with NIRS data alone had the highest prediction ability in only 1 of 28 scenarios for within-environment prediction. Potential causes of the surprisingly lower phenomic than genomic prediction power in this study are discussed, including sample size, sample homogenization, and low G×E. A genome-wide association study (GWAS) implicated known (i.e., *MADS69*, *ZCN8, sh1, wx1, du1*) and unknown candidate genes linked to plant height and flowering-related agronomic traits as well as compositional traits such as kernel protein and starch content. This study demonstrated that including NIRS with genomic markers is a viable method to predict multiple complex traits with improved predictive ability and elucidate underlying biological causes.

**Key message:** Genomic and NIRS data from a maize diversity panel were used for prediction of agronomic and kernel composition traits while uncovering candidate genes for kernel protein and starch content.

## Introduction

Historically, grain yield improvement has been the primary objective of many public breeding programs. However, depending on germplasm and environments screened, there are often strong phenological associations between yield and kernel-, height-, and flowering-related traits in *Zea mays* L. (maize, or corn) (Adak et al. 2021b; Adak et al. 2021c; Baye et al. 2022; Boomsma et al. 2010; Borrás et al. 2004; Dai et al. 2021; Farfan et al. 2013; Pedersen et al. 2022; Ribaut et al. 1996; Westgate and Boyer 1986). Currently in commodity markets, farmers are only compensated for grain yield (meeting quality thresholds) in commercial maize production, but the development of specialty niche markets requiring higher quality grain is made on contract for specific uses, such as whiskey distillation (Arnold et al. 2019).

In terms of agronomic traits related to maize grain yield, plant height is a highly heritable trait with many characterized large-effect alleles relevant for overall fitness and agricultural performance (Andorf et al. 2010; Fernandez et al. 2009). In the U.S. Corn Belt where plants are tall owing to optimal growing conditions, taller plants lodge more frequently so height can be negatively correlated to yield; in Texas and the south where lodging is rarer, taller plants tend to yield more (Farfan et al. 2013). Therefore, breeding efforts for yield have indirectly selected for height adaptations in monocultured maize (Peiffer et al. 2014). Characterizing flowering time is also of interest to breeding programs given it represents a plant’s adaptation to its environment. Changes in flowering time serve as a strategy to avoid abiotic (e.g., heat) and biotic stresses and mature grain before winter (Buckler et al. 2009; Song et al. 2017). Yield and flowering can be interlinked, for example with water stress during anthesis affecting floral development, while longer anthesis-silking intervals (ASI) are associated with decreased grain yield (Ribaut et al. 1996; Westgate and Boyer 1986). Kernel composition traits such as phosphorus content are of interest due to their implications for human and animal health, as well as in nutrient loading of waterways (Lorenz et al. 2007; Raboy 2002; Raboy et al. 1989; Wardyn and Russell 2004). Other kernel composition traits such as protein and starch are of interest to breeders and geneticists and are directly related to grain yield of cereal crops, with protein negatively correlated to yield and starch positively correlated (Berke and Rocheford 1995; Goldman et al. 1993; Prioul et al. 1999; Séne et al. 2000; Wilson et al. 2004).

Near infrared reflectance (NIR) spectroscopy (NIRS) is a remote sensing approach that leverages techniques in spectroscopy, analytical chemistry, and mathematics (specifically chemometrics) (Pasquini 2018). NIRS functions by barraging a sample with electromagnetic radiation at wavelengths typically between 750 to 2500 nm, producing absorption or reflectance values of any compound containing C-H, N-H, S-H, or O-H chemical bonds (Pasquini 2003, 2018). NIRS measurements are generally recorded in laboratory settings using dried tissue or grain (Ferrio et al. 2004), although portable spectrometers can provide non-destructive measurements in the field. NIRS machines integrate each sample to a single point measurement and differs from hyperspectral NIR imaging which creates an image but measures fewer discrete bands. Hernandez et al. (2015) demonstrated the ability to predict grain yield using NIRS measurements collected at anthesis and grain fill as a proxy trait for grain yield in a diverse wheat panel. Recently, some small plot combines and forage harvesters have been equipped with NIRS to minimize labor and bias of subsampling grain or biomass (Polytec GmbH 2022). Given the prevalence of NIRS in plant science and breeding programs for composition and quality testing, many researchers are already equipped to incorporate NIRS as a method for phenomic prediction using reflectance values as markers instead of genomic markers, in turn facilitating phenomic selection (PS). In the first major phenomic prediction study, Rincent et al. (2018) demonstrated the ability of NIR spectra to enable prediction accuracy of complex traits on par with marker-based selection, despite subjecting the models to environments distinct from the one in which NIRS data were recorded. The study employed covariance matrix estimation in the manner of Yamada et al. (1988) to predict wheat yield in a separate environment, highlighting the role NIRS relationship matrices can play in demarcating genetic relationships between lines. NIR spectra have been used previously to estimate complex plant compositional traits and used to predict grain yield based on the relationships that exist between grain yield and composition. Spectra can also be implemented in combination with molecular markers to bolster genomic prediction accuracy or coupled with a more commonly recognized genomic relationship (kinship) matrix (Robert et al. 2022b). Omics-based selection was proposed on the basis that phenomic markers, such as those afforded by NIRS, capture genetic relatedness, enabling cross-environment predictions from spectra obtained in different environments. This concept, demonstrated in Rincent et al. (2018), produced predictions with higher accuracy at times than GS (Robert et al. 2022b). Weiß et al. (2022) reported more stable prediction ability of NIRS phenomic vs. genomic prediction when predicting dry grain matter content, grain yield, and phosphorus concentration across diverse breeding material. Weiß et al. (2022) also found genomic prediction to be less tolerant to genetic divergence between training and test data sets. Robert et al. (2022a) demonstrated the combined predictive power of genotypic and NIRS data and emphasized the impact of environment on predictive ability as captured by NIRS using two years of winter bread wheat candidate breeding lines. In a large panel of soybean recombinant inbred lines, Zhu et al. (2021a) found phenomic prediction ability to be comparable, and in some cases greater than, genomic prediction ability within and between segregating families. Zhu et al. (2021a) also found that NIRS data sets from distinct environments can be substituted for data sets from other environments and applied successfully in phenomic prediction. This is potentially attributable to phenomic selection associating endophenotypes with target traits, meaning the association itself is required for accurate prediction, allowing for variation in the endophenotypes captured between environments (Zhu et al. 2021a).

In this study, building on NIRS data from Lane et al. (2020) and genomic data from Farfan et al. (2015), phenomic (NIRS) and genomic data of these maize lines evaluated as hybrids on a common tester were used as predictors either separately or together to predict seven agronomic traits and three kernel composition traits. To assess prediction accuracy in real plant breeding scenarios, the performance of tested and untested maize hybrids was predicted in observed and unobserved environments across water stressed and well-watered growth conditions over two years (four environments total) resulting in four distinct cross-validation schemes. Within-environment prediction models were also evaluated in each of the four environments using a single cross-validation scheme. In summary, this research sought to: i) develop combined genomic and NIRS models capable of predicting traits such as grain yield and protein content in maize across and within environments; ii) assess the utility of including NIR spectra in predictive models alone and alongside genotypic data; and iii) to use this data to uncover candidate genes pertaining to agronomically relevant maize traits across four distinct growing environments to shed light on the genetic basis of these phenomic and genomic predictions.

## Materials and Methods

### Experimental Design

Field trials were conducted at the Texas A&M AgriLife Experiment Station in Burleson County, Texas, during the summers of 2011 and 2012 as described in detail in Farfan et al. (2015). Briefly, a subset of 346 maize lines from the 302-line USDA Goodman maize association panel (Flint-Garcia et al. 2005) and the 300-line diverse southern subtropical Williams/Warburton panel (Warburton et al. 2013) were testcrossed to two variants of Tx714 (Betrán et al. 2004). These two variants of Tx714, isogenic lines with lipoxygenase 4 and 5 knockouts, were not significantly different for any measured trait and were therefore combined throughout analysis. Testcross hybrids were grown in randomized complete block designs with two replications in each year and management. Experimental units were one-row plots that were 7.92 m long and 0.76 m wide with a target plant density of 75,000 plants/ha. Plots were planted according to a range/row design where ranges refer to horizontal gridlines running perpendicular to tractor rows and rows refer to vertical gridlines parallel with tractor rows. Testcross hybrids were grown in College Station, TX, USA (CS) in a Ships clay loam soil and subjected to well-watered (WW) and water stressed (WS) management conditions in 2011 and 2012. These will hereafter be denoted as CS11_WS, CS11_WW, CS12_WS, and CS12_WW indicating four different growing environments. CS11_WS, CS11_WW, CS12_WS and CS12_WW contained 199, 270, 319 and 220 maize hybrids, respectively, of which a subset of 145 was common to all four environments.

### Agronomic Traits

Seven agronomic traits related to yield, flowering, and height were recorded on a row-plot basis. Additionally, three kernel composition traits (phosphorus, protein, and starch) were estimated from NIRS scans, however results of genomic, phenomic, and combined genomic/phenomic prediction for these traits are reported as supplementary figures due to the potentially confounding nature of using phenomic data to predict trait values derived from estimates based on the same data (as opposed to all values obtained from wet chemical analysis). **Table 1** describes each trait and its associated abbreviation, unit, and trait category. The kernel composition traits were calculated by scanning the maize kernels in a Thermo Scientific Antaris II Fourier transformed interferometer (Thermo Fisher Scientific, Waltham, MA, USA) and applying calibrations reported in Meng et al. (2015) and Christman (2017). Due to extreme drought conditions in the CS11_WS environment, data were not recorded for kernel composition traits in the water stressed environment which led to atypically low yield. Data for all other traits were obtained for each of the four environments (**Fig. 1b**).

**Figure 1.**
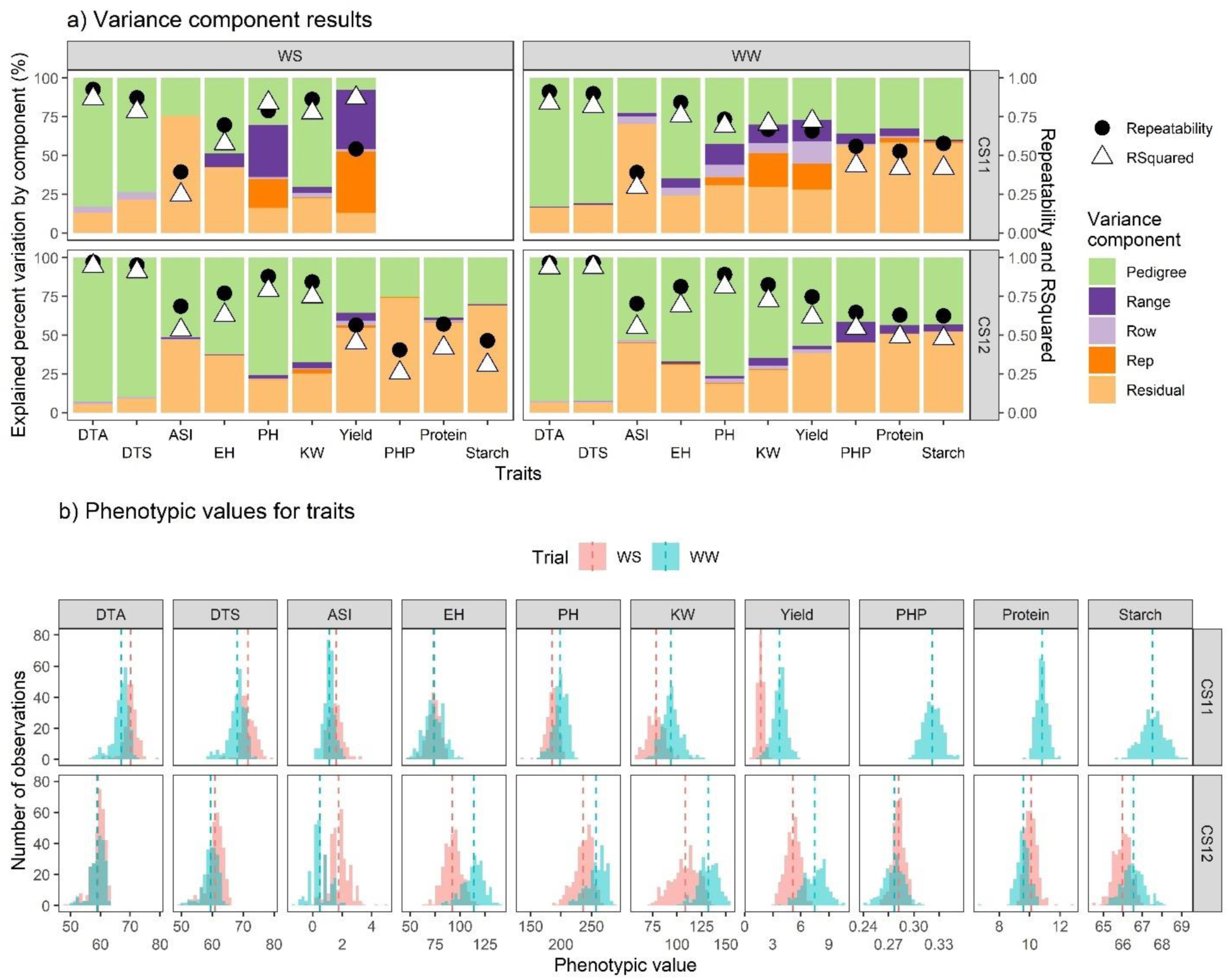
(**a**) Results are displayed for ANOVA of agronomic traits according to **Eq. 1**. Black circles denote repeatability and white triangles indicate R^2^ values. (**b**) Phenotypic data distributions are shown for each trait with red denoting water stress and blue denoting well-watered conditions. Dashed lines indicate the means of each distribution. X-axes differ according to the units of the reported agronomic traits (as given in **Table 1**). Phenotypes are not reported for protein, PHP, and starch in CS11_WS due to atypical heat and drought that severely affected yield in this environment.

**Table 1.** The seven agronomic traits of interest are presented alongside their abbreviations, units in which they were recorded, and the overall category they belong to. Three kernel composition traits estimated from NIRS scans are also presented. DAP refers to days after planting.

| **Trait** | **Abbreviation** | **Units** | **Category** |
| --- | --- | --- | --- |
| Days to anthesis | DTA | DAP of 50% anther production in plot | Flowering |
| Days to silking | DTS | DAP of 50% silk production in plot |  |
| Anthesis-silking interval | ASI | Days between silking and anthesis |  |
| Ear height | EH | Height in cm from ground to shank of first ear | Plant height |
| Terminal plant height | PH | Height in cm from ground to tip of tassel |  |
| Kernel weight | KW | 500 kernels (g) | Yield |
| Grain yield | Yield | t/ha |  |
| Phosphorus | PHP | Percent of dry matter | Kernel composition |
| Protein | N/A | Percent of dry matter |  |
| Starch | N/A | Percent of dry matter |  |

### Phenomic Data: Near Infrared Spectroscopy

NIRS data from grain harvested from the testcross hybrids developed by Farfan et al. (2015), as reported in Lane et al. (2020) were used. In brief, whole kernel samples of 199 (CS11_WS), 270 (CS11_WW), 319 (CS12_WS) and 220 (CS12_WW) hybrids were scanned in a rotating cup using a Thermo Scientific Antaris II Fourier transformed interferometer. Scanning and spectral processing methodologies are described in Meng et al. (2015) and initial analyses and quality control parameters were discussed in Lane et al. (2021). As a result, 3112 NIRS bands were extracted from each sample. A brief overview of the scanning methodology is as follows: a) whole kernel samples in ∼175 g batches were analyzed in triplicate with 128 scans per sample; b) with the scanner set to record diffuse reflectance, scans were performed over the instrument’s integrating sphere module; c) all spectra were recorded at ambient temperature and were calculated at 4 cm^-1^ resolution, thus encompassing 4,000-10,000 cm^-1^; d) preprocessing techniques such as multiplicative signal correction, standard normal variate, data treatment with first and second derivative transformations, as well as Savitzky-Golay filtering was performed in Thermo Scientific’s TQ Analyst Pro Edition Software (Meng et al. 2015).

Using the *caret* package in R (Kuhn 2008), the Lasso algorithm was run with 5-fold-3-replication cross-validation using the maximum available hybrids in each environment to determine the most important NIRS bands (cm^-1^) associated with grain yield and kernel composition traits. Model performance was assessed using R^2^ (**Supplementary Data 1**).

### Genotypic Data

Using the Genotype By Sequence (GBS) method outlined in Elshire et al. (2011), genome-wide single nucleotide polymorphism (SNP) genotypic data were obtained for 213 lines from the Williams/Warburton panel as performed by the USDA-ARS Corn Host Plant Resistance Research Unit (Mississippi State University, MS). Publicly available data in Panzea (Zhao et al. 2006), genotypic information was obtained from 133 lines of the 282 available inbred lines comprising the USDA Goodman maize panel which used the same genotyping method. These GBS data were used for the genome-wide association study (GWAS), genomic, and combined genomic/phenomic prediction models. SNPs were filtered from an initial array of 61,410 polymorphic loci, and SNPs with a MAF < 0.05 were discarded.

Filtering of genomic data was performed as follows: (i) heterozygote calls were set as missing data; (ii) markers with greater than 20% missing values were removed; (iii) markers with less than 0.05 minor allele frequency were removed. After filtering, 34,145 markers (according to maize genome version AGPV2) were used in the GWAS. An all-random model was used to calculate the explained percent variation of significant loci in GAPIT (Lipka et al. 2012). Two Bonferroni-adjusted significance thresholds were set based on alpha (α) values of 0.05 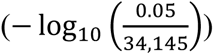 and 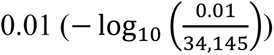, resulting in cutoffs of 5.8 and 6.5 respectively (**Fig. 7**).

### Analysis of variance

To estimate genetic and other experimental variance components, **Eq. 1** was applied to the seven agronomic traits, three kernel composition traits estimated via NIRS, and each NIRS band (3112 bands) separately in each of the four environments (CS11_WS, CS11_WW, CS12_WS, and CS12_WW) to obtain the best linear unbiased predictor (BLUP) additive genetic effects via the *lme4* package in R (Bates et al. 2014).

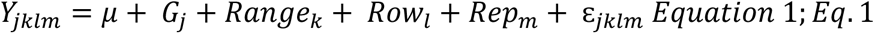

*Y* is a vector denoting each of the seven agronomic traits, kernel composition traits, or each of the NIRS bands for observations of the *j*th maize hybrid in the *k*^th^ range, *l*^th^ row, and *m*^th^ replication; *μ* denotes the grand mean; *G* denotes the effect of the *j*^th^ maize hybrid where 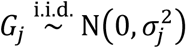; *Range* denotes the effect of the *k*^th^ range where *Range*_*k*_ 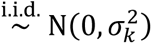; *Row* denotes the effect of *l*^th^ row where *Row*_*l*_ 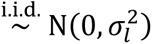; *Rep* denotes the effect of the *m*^th^ replication where *Rep*_*m*_ 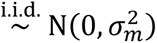; and lastly, ɛ denotes the combined error (σ^2^*_error_*) of the residuals of the variance components of the model where 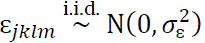.

Individual environmental repeatability was calculated using **Eq. 2**.

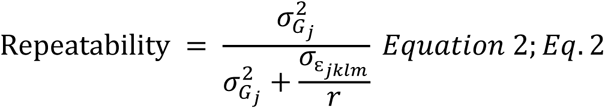

Here, 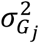, and 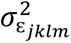 are the variance of hybrids and error respectively, and *r* is the number of replications. The genetic effects for NIRS bands across each of the four environments and for seven agronomic traits are viewable in **Supplementary Data 1**.

### Across-environment genomic and phenomic predictions

For across-environment prediction schemes, the 145 maize hybrids common to all four environments were considered. The *BGLR* package was used to perform the predictions of seven maize agronomic traits and three kernel composition traits in R (Pérez and de Los Campos 2014) considering genomic and/or phenomic covariates. Models were fit under a hierarchical Bayesian framework, with 5000 Gibbs sampler iterations, 20% burn-in, and the thinning parameter was set at 10 for a total of 400 iterations used to compute the posterior mode of the different parameters involved in the linear predictors.

Three genomic best linear unbiased prediction (GBLUP) models based on covariance structures were compared in this study (**Table 2**). The data preparation and matrix multiplication to obtain each kernel is described at (https://bit.ly/Maize-NIR-GBS). In the series of equations, *Y* represents a vector of BLUPs of seven agronomic traits and three kernel composition traits for each of the 145 hybrids in four environments; *μ* represents the grand mean; *E* represents the environment matrix (CS11_WS, CS11_WW, CS12_WS, and CS12_WW) which was modelled as: 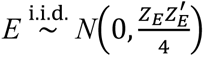, where *Z*_*E*_ is the block diagonal incidence matrix.

**Table 2.**
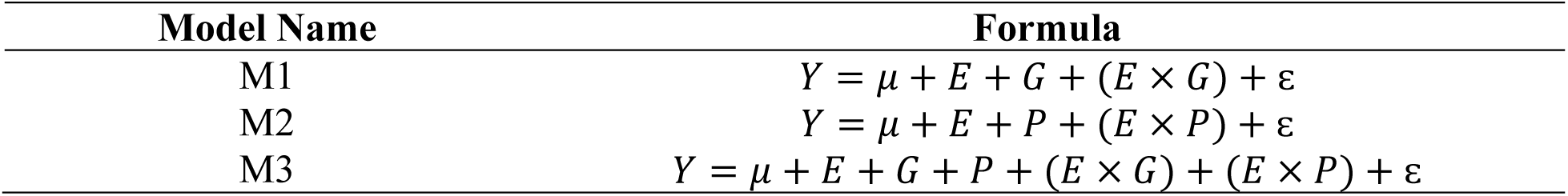
Summary of prediction models for across-environment predictions. M1 includes the environment (E), additive genomic (G), and E×G kernels, M2 contains the E, NIRS phenomic (P), and E×P kernels, and M3 contains E, G, P, E×G, and E×P kernels.

| Model Name | Formula |
| --- | --- |
| M1 | $Y = \mu + E + G + (E \times G) + \varepsilon$ |
| M2 | $Y = \mu + E + P + (E \times P) + \varepsilon$ |
| M3 | $Y = \mu + E + G + P + (E \times G) + (E \times P) + \varepsilon$ |

*Genomic data preparation* – *G* symbolizes the additive genomic effects, modeled via a linear combination between *n* additive markers and their effects based on VanRaden (2008) as follows: 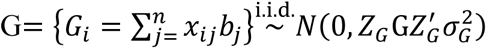 where 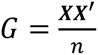, *G* is the column-wise centered and standardized matrix of additive markers of inbreds that were topcrossed with Tx714, and 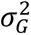 denotes the corresponding variance components. Because only one pollinator (Tx714) was used in the topcross, only an additive genomic relationship matrix was calculated. The additive genomic relationship matrix was generated according to VanRaden (2008) using the *AGHmatrix* package in R (Amadeu et al. 2016).

*Phenomic data preparation* – The phenomic effect (*P*) associated with matrix of NIRS bands (*N*), similar to the genomic effect matrix, was modelled via a linear combination of *p* NIRS bands and their effects as follows: 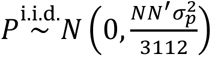 where 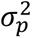 is the associated variance component. The interaction terms between environment and additive effects or phenomic (NIRS) effects were modeled corresponding to (Jarquín et al. 2014). Briefly, the Hadamard product (where ⊙ denotes the element-wise product) of the covariance structure of the interaction terms was modeled as follows: 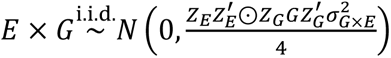 and 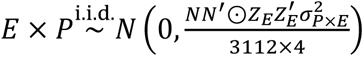, where 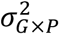 and 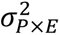 are the corresponding variance components. Environment effects were calculated using Bayesian ridge regression, and all other kernels were specified as Reproducing Kernel Hilbert Spaces Regressions (specified as “BRR” and “RKHS” respectively in the *BGLR* package documentation) which corresponds to the GBLUP fashion of the model terms. *Z_G_* and *Z_E_* are the incidence matrices that connect phenotypes with genotypes and environments, respectively.

### Within-environment prediction

In within-environment prediction, 199, 270, 319 and 220 hybrids were used in CS11_WS, CS11_WW, CS12_WS, and CS12_WW respectively (the maximum possible for each environment). Three prediction models were compared in the within-environment prediction framework. The genomic and phenomic NIRS relationship matrices were obtained in the same manner as described for across-environment prediction (**Table 3**). Only main effects were considered due to predictions occurring strictly within environments.

**Table 3.** Summary of prediction models for within-environment predictions. M1 specifies a model that incorporates only genomic effects; M2 specifies a model that incorporates only NIRS (phenomic) effects; M3 specifies a model that incorporates both genomic and NIRS effects.

| Model Name | Formula |
| --- | --- |
| M1 | $Y = \mu + G + \varepsilon$ |
| M2 | $Y = \mu + P + \varepsilon$ |
| M3 | $Y = \mu + G + P + \varepsilon$ |

### Cross-validation performance evaluation

The prediction performance of each model was assessed using a 5-fold-25-replication cross-validation scheme in both across– and within-environment predictions. In across-environment prediction, four prediction scenarios were implemented based on the naming convention proposed by Jarquín et al. (2017), where CV2 represents prediction of tested genotypes in observed environments, CV1 represents prediction of untested genotypes in observed environments, CV0 represents prediction of tested genotypes in unobserved environments, and lastly CV00 represents prediction of untested genotypes in unobserved environments. In CV1 and CV2, training data consisted of genotypes belonging to all four environments, so all four environments were considered tested environments. In CV2 and CV1, the predictive ability was assessed as the correlation between predicted and observed phenotypic records within the same environment.

For the within-environments or single environment prediction, only CV1 was calculated for and is denoted CV1-W. An illustration of the cross-validation schemes is presented in **Supp.** Fig. 1.

In CV0 and CV00, models were trained using the information of tested genotypes belonging to only well-watered environments in 2011 and 2012 (CS11_WW and CS12_WW), so water stressed environments in 2011 and 2012 (CS11_WS and CS12_WS) were considered untested environments (the training data set size was reduced by half and restricted to CS11_WW and CS12_WW). Simply, the well-watered environments served as training data and water stressed environments served as test (validation) data. In CV0, prediction ability was calculated as the correlation between actual and predicted values of tested genotypes across environments, and in CV00 prediction ability was assessed as the correlation between actual and predicted values of untested genotypes across environments (**Supp.** Fig. 1).

### Prediction Ability

To quantify prediction ability of each CV/model/trait combination, a weighted average correlation was implemented according to Tiezzi et al. (2017) for across-environment predictions as described in **Eq. 3**. The same implementation of prediction ability was used for within-environment predictions. Prediction abilities are grouped by fold and replication and calculations of the individual components of this equation are given in **Supplementary Data 1**.

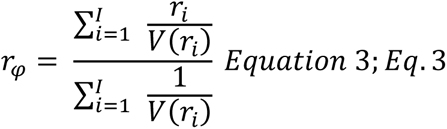

Where *r*_*i*_ is the Pearson’s correlation between actual and predicted values belonging to the *i*^th^ environment, 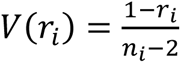 is the sampling variance, and *n* is the number of hybrids in each environment.

### Genome-wide association study

To discover loci linked to the seven agronomic traits collected in each of four environments, a genome-wide association study (GWAS) was run with the BLINK method (Huang et al. (2019) using the *GAPIT* (Version 3) package in R (Lipka et al. 2012). 199, 270, 319, and 220 hybrids were used in GWAS in CS11_WS, CS11_WW, CS12_WS and CS12_WW respectively. Filtering of genomic data was performed as described previously in the *Genomic data preparation* segment.

All statistical analyses were performed using R Version 4.2.2 within the RStudio application Build 353 (R Core Team, 2022; Posit Software, 2023). An annotated script of the code used in this study is available on GitHub (https://github.com/ajdesalvio/Maize-NIRS-GBS.git). Files necessary to run the script are available within this repository.

## Results

### Variance component analysis for agronomic traits

Notable fluctuations in variance components were observed across traits and environments (**Fig. 1)**. A large proportion of the total variation for DTA and DTS was explained by pedigree (74 to 93 percent) which led to high repeatability values (0.87 to 0.97). However, only 22 to 53 percent of the total variation was explained by pedigree for ASI, which had a lower range of variation (**Fig. 1b**), leading to lower (0.25 to 0.53) repeatability. For height-related traits, pedigree explained between 30 to 76 percent of the total variation for PH and EH that produced repeatability values of 0.87 to 0.97. Within yield-related traits, pedigree explained between 10 to 70 percent of the total variation for yield and KW while repeatability ranged from 0.54 to 0.86. For the kernel composition traits, pedigree explained between 25 to 43 percent of total variation for phosphorus (PHP), protein, and starch content, and repeatability values were between 0.40 to 0.65 (**Fig. 1a**).

Distributions of maize phenotypic data show distinctions between water stressed and well-watered environments across all traits in one or both years (**Fig. 1b**). These differences in phenotype distributions were consistent in direction between environments within each year. For instance, ASI values were greater in water stressed environments than well-watered environments in both years. Conversely, KW and yield values were greater in well-watered environments than water stressed environments in both years (**Fig. 1b**).

### Variance component analysis for NIRS bands

The results of ANOVA using **Eq. 1** across all 3112 NIRS wavelength bands revealed trends of explained percent experimental variation across each band that were similar in each of the four growing environments (**Fig. 2a**); however, overall patterns of each band’s respective variance components were responsive to each environment. On average, genotypic variance (pedigree) explained the highest percentage of experimental variation in all environments, fluctuating between 32 and 79 percent among NIRS bands. Depending on the environment, the highest explained percent variation by pedigree was in CS11_WS (between 49 and 78 percent), which also had the larger range of variation (**Fig. 2b**). Repeatability, as computed by **Eq. 2**, ranged between 0.50 and 0.88. The highest overall values for repeatability were observed in CS11_WS. In addition, repeatability was found to increase from ∼0.50 to ∼0.85 in each environment when observing the NIRS bands increase from 4,000 to 10,000 cm^-1^. Phenotypic values of NIRS bands over the entire spectrum followed similar trends in each environment (**Fig. 2b**). Field spatial variation (range, row, and replication effects) was most pronounced in CS11_WW, similar to phenotypic traits (**Fig. 1a**), though it remained at or below 10 percent across the entire spectrum.

**Figure 2.**
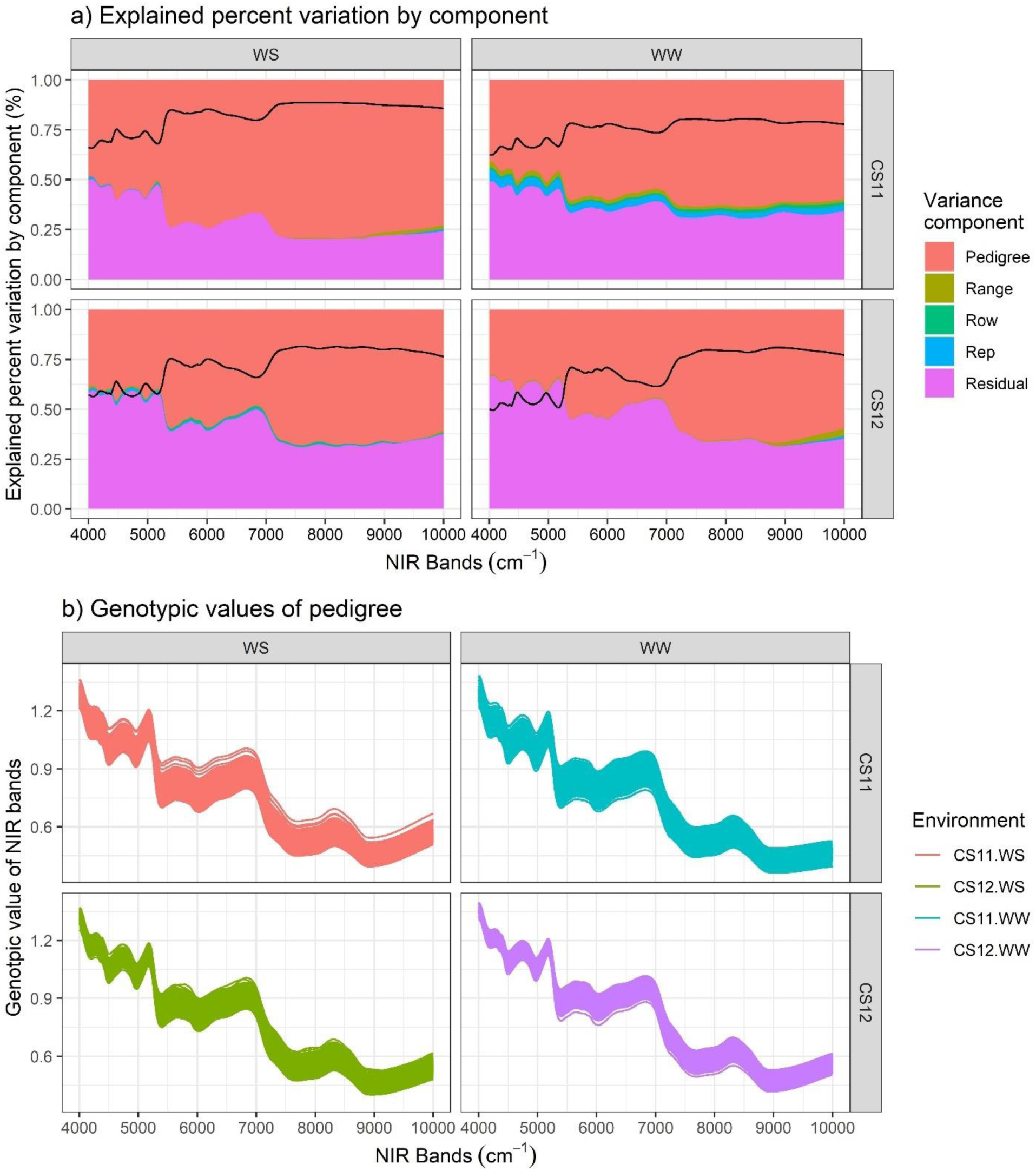
(**a**) Explained percent variation across all 3112 NIRS bands in each of the four environments. 2a is displayed as a proportional stacked area chart. Black lines in each of the four environments denote repeatability values as they change across the bands (**Eq. 2**). (**b**) Genotypic values of all NIRS bands are displayed for each environment as calculated in **Eq. 1**.

### Explained percent variation by phenomic and genomic prediction models

#### Across-environment

The environmental component (*E*) commonly explained the highest percent of experimental variation depending on the prediction model and trait reaching up to 89% of total variation (**Fig. 3a**). For yield-related traits, in M3 the phenomic effect (*P*=46%) explained variation of KW greater than genomic effects (*G*=4%), and combined, these effects surpassed the environment effect (40%). Similarly for yield, phenomic effects (15%) explained greater variation than genomic effects (2%) in M3. The G×E kernel (*EG*) and the NIRS×E kernel (*EP*) explained a minor amount of experimental variation in each of the seven agronomic traits. For genomic prediction (M1) of kernel composition traits, the environment component (*E*) explained the highest percentage of experimental variation for all three traits, with as much as 82% for PHP (**Supp.** Fig. 2a).

**Figure 3.**
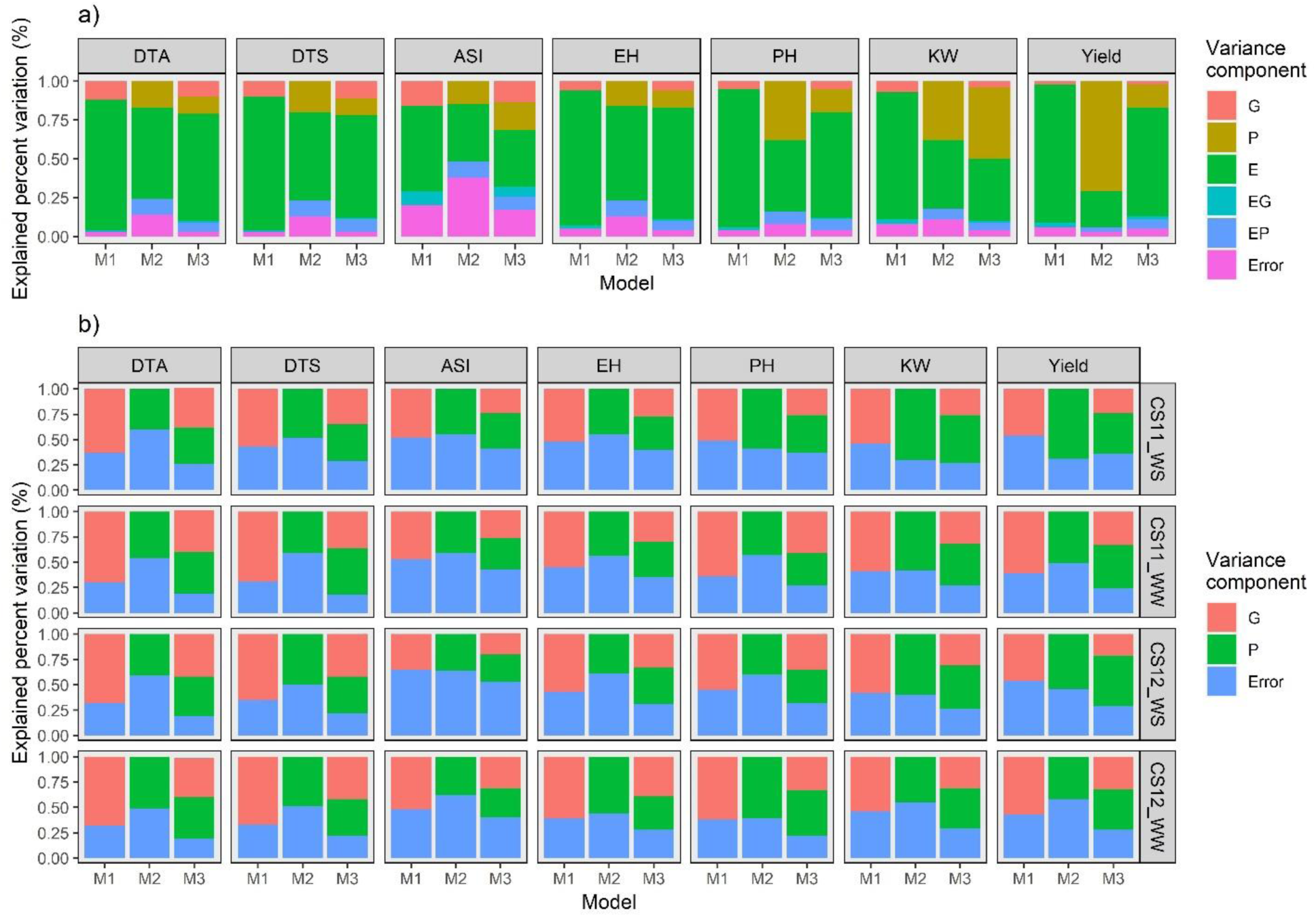
Explained percent variation of genomic/phenomic prediction model components is displayed for each model across all agronomic traits. Variance component breakdowns are reported for each model for (**a**) across-environment prediction and (**b**) within-environment prediction, where G: genomic kernel; P: NIRS (phenomic) kernel; E: environment kernel; EG: interaction kernel between genomic and environment kernels; EP: interaction kernel between NIRS (phenomic) and environment kernels; Error: unexplained (residual) variation. M1: E, G, and EG prediction kernels; M2: E, P, and EP prediction kernels; M3: E, G, P, EG, and EP prediction kernels.

#### Within-environment

In general, genomic effects in M1 explained a greater percentage of experimental variation than phenomic effects in M2 for agronomic traits across four environments **(Fig. 3B**). The explained percent variation by genomic and phenomic effects in M1 and M2 changed between 35% to 70% (**Fig. 3B**). The explained percent variation by error was reduced in M3 compared to M1 and M2 models. For yield-related traits predicted by M3, phenomic effects consistently explained greater variation than genomic effects. For genomic prediction (M1) of the three kernel composition traits, the genomic kernel always explained a greater percentage of experimental variation than the residual, ranging between 51% to 57% (**Supp.** Fig. 2b).

### Assessment of Prediction Ability: Across-Environment Prediction

Each of the cross-validation schemes (CV00, CV0, CV1, CV2) that comprised various simulated plant breeding scenarios (described in detail by Crossa et al. (2017) and implemented in Jarquín et al. (2021)) were adapted for this study with the aim of using training data from well-watered environments to predict phenotypic values of agronomic traits in water stress environments. A summary of model performance and standard deviations for each CV/model/trait combination is presented in **Fig. 4** and results are viewable within the supplementary data repository.

**Figure 4.**
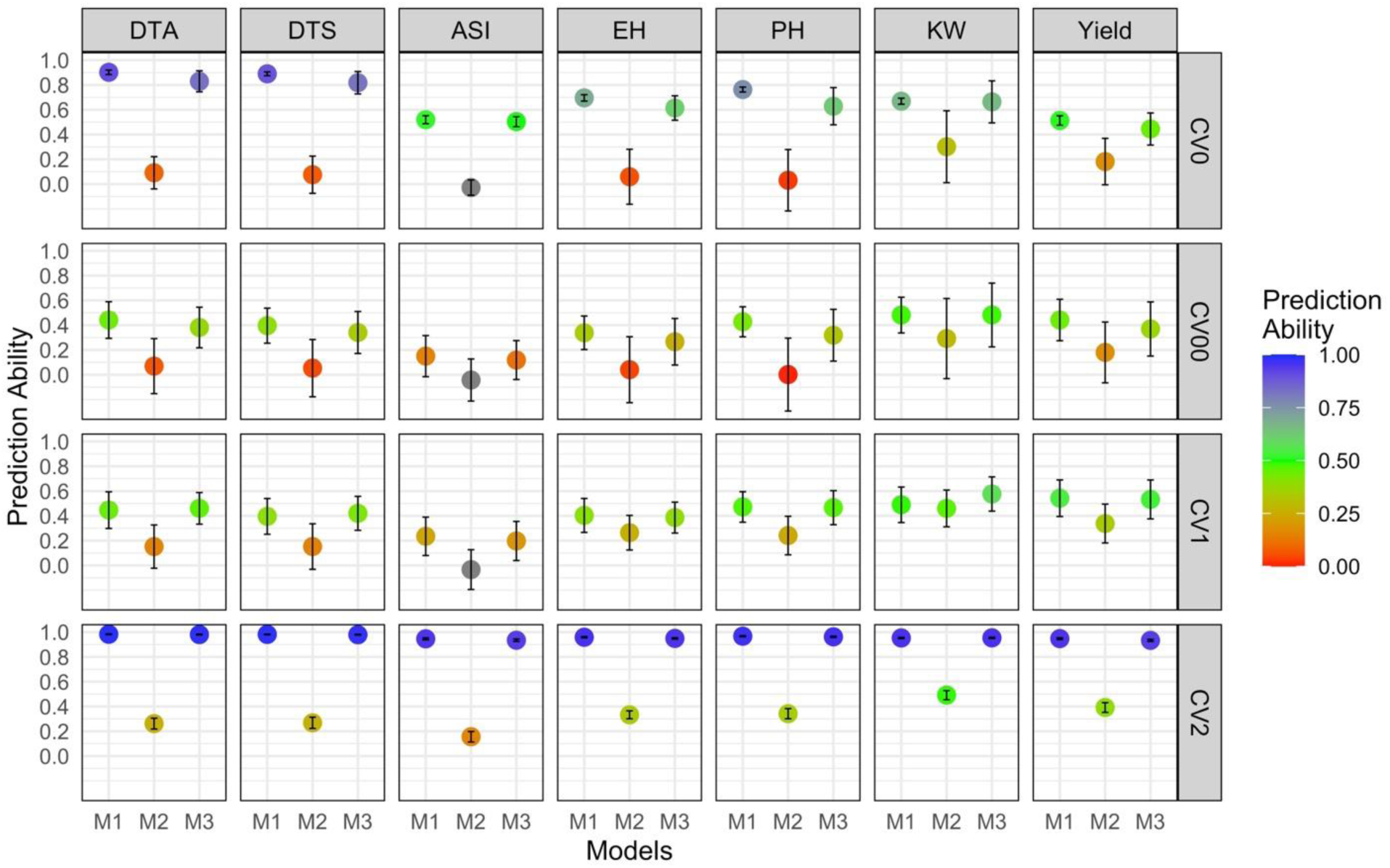
Prediction abilities of each across-environment prediction model for all traits are displayed stratified by trait and cross-validation scheme. Prediction abilities close to 1 are indicated by a dark blue color while those closer to 0 appear red. Model notation is described in detail in **Table 2**. M1: environment (E), genomic (G), and environment×genomic (EG) prediction kernels; M2: E, phenomic (P), and environment×phenomic (EP) prediction kernels; M3: E, G, P, EG, and EP prediction kernels.

Average prediction abilities for all models grouped by CV were 0.38±0.16 for CV00, 0.71±0.25 for CV0, 0.44±0.14 for CV1, and 0.96±0.0047 for CV2 (**Fig. 4**). For prediction of untested genotypes in uncharacterized environments (CV00), the most difficult prediction scheme across environments, the genomic and environmental effect model (M1) had the highest prediction ability in six of the seven agronomic traits, with 0.44±0.15 for DTA, 0.40±0.14 for DTS, 0.15±0.17 for ASI, 0.34±0.14 for EH, 0.43±0.12 for PH, and 0.44±0.17 for yield. The combined genomic/phenomic and environment model (M3) performed best for KW (0.48±0.26), marginally outperforming M1. Genomic prediction (M1) outperformed the other models for all seven traits when predicting tested genotypes in uncharacterized environments (CV0). For prediction of untested genotypes in characterized environments (CV1), M1 displayed the highest prediction ability for ASI, EH, PH, and yield, while M3 outperformed the other models for predicting DTA, DTS, and KW. For prediction of tested genotypes in characterized environments (CV2), M1 outperformed all other models (only slightly beating M3) for prediction of six of the seven agronomic traits excluding KW, which was again best predicted by M3 by a small margin (**Fig. 4**). For across-environment genomic prediction (M1) of kernel composition traits (PHP, protein, and starch), average prediction abilities according to CV were 0.11±0.16 for CV00, 0.22±0.05 for CV0, 0.38±0.16 for CV1, and 0.95±0.0057 for CV2 (**Supp.** Fig. 3).

### Assessment of Prediction Ability: Within-Environment Prediction

In the context of within-environment prediction, only one cross validation scheme can be usefully evaluated. Here, CV1-W refers to the prediction of untested genotypes in a single characterized environment (as opposed to all four). A summary of the performance of models for each model/trait/environment combination is presented in **Fig. 5** and **Supplementary Data 1**. In nearly every prediction scenario, the performances of M1 and M3 were similar. DTA was best predicted by the combined genomic and phenomic model (M3) and slightly outperformed M1 in all environments (average prediction ability of 0.65±0.06) except for CS11_WS, where the genomic prediction model (M1) had the highest prediction ability (0.52±0.08). DTS was best predicted again by M3 in three environments (average prediction ability of 0.58±0.07) except for CS11_WW, where M1 had a prediction ability of 0.61±0.10. Prediction ability for ASI was low, with M1 outperforming other models (averaging 0.15±0.13) except in CS12_WS, where the phenomic prediction model (M2) performed best (0.15±0.16), the only instance of a phenomic-only model (M2) outperforming M1 and M3 for within-environment prediction. EH was mostly predicted best by M1 (average 0.52±0.10) except in CS11_WS, where the combined model (M3) performed slightly higher than the others (0.40±0.26). M3 had the highest prediction ability for all four environments for KW (average 0.53±0.11). PH and yield were evenly split between M1 and M3 having the highest prediction abilities, with M1 averaging 0.56±0.07 and 0.58±0.15 for PH and yield respectively, and M3 averaging 0.55±0.10 for PH and 0.53±0.09 for yield across the four environments (**Fig. 5**). For within-environment genomic prediction (M1) of kernel composition traits (PHP, protein, and starch), prediction abilities were moderate regardless of the trait or environment, averaging 0.30±0.14, 0.29±0.13, and 0.34±0.17 for PHP, protein, and starch respectively (**Supp.** Fig. 4).

**Figure 5.**
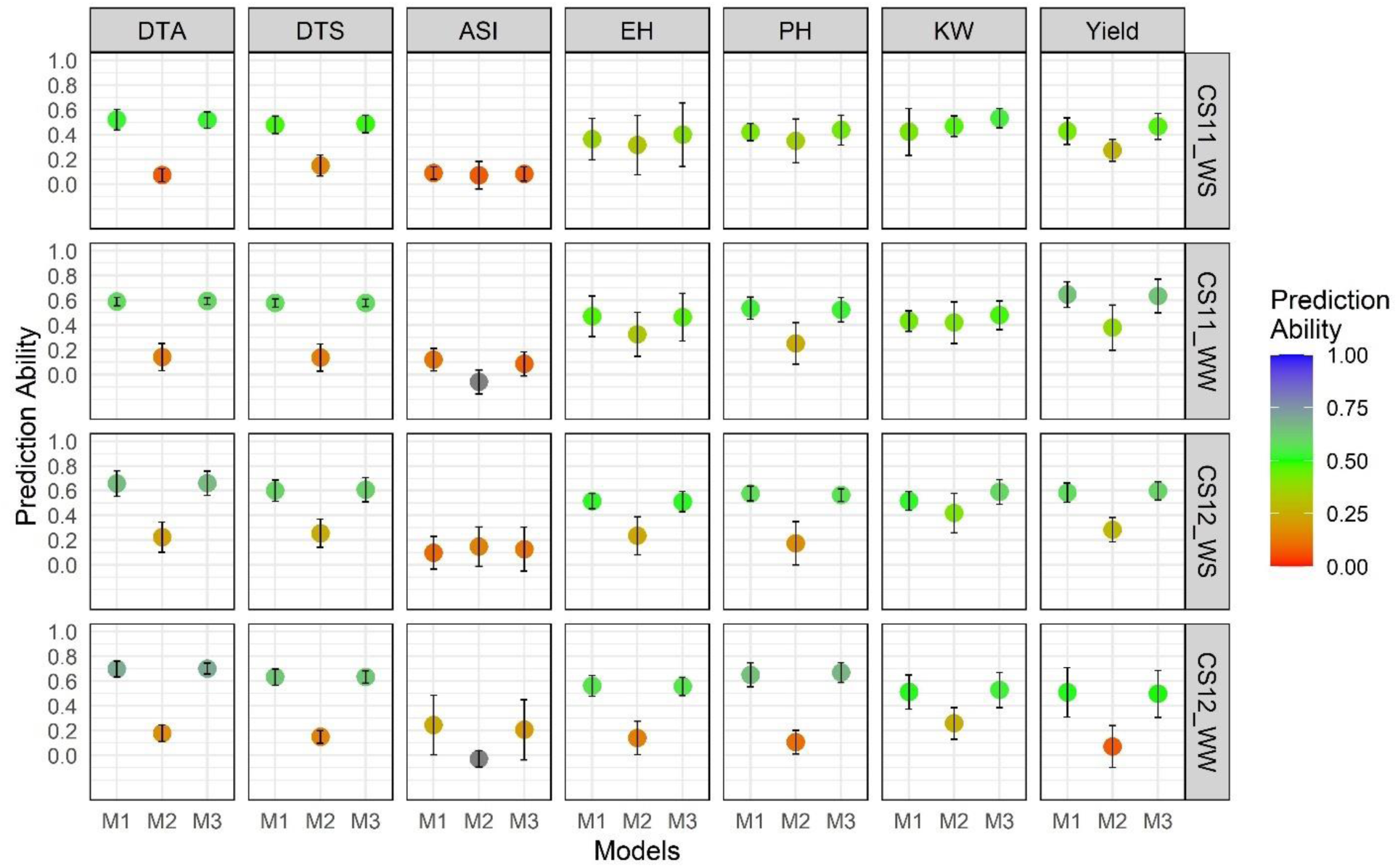
Prediction abilities of each within-environment prediction model for all traits are displayed stratified by trait and environment. Only one cross-validation scheme was implemented for within-environment prediction (prediction of unknown genotypes in characterized environments, CV1-W). Prediction abilities close to 1 are indicated by a dark blue color while those closer to 0 appear red. Model notation is described in **Table 3**. M1: genomic (G) prediction kernel; M2: phenomic (P) prediction kernel; M3: G and P prediction kernels.

### Correlations within and between agronomic traits and NIRS bands across environments

Correlations between height– and flowering-related (except for ASI) traits were consistently positive (ranging between 0.43 to 0.70) across four environments. Similarly, correlations between height– and yield-related traits were consistently positive (ranging between 0.15 to 0.57) across four environments. However, correlation values between flowering– and yield-related (except for ASI) traits were not consistent (ranging from –0.23 to 0.41) across four environments (**Fig. 6a**). Kernel composition traits (except for starch) generally had negative correlations with yield-related traits (ranging from –0.42 to 0.07). Correlations of the 3112 NIRS band wavelengths with each other revealed almost every pairwise correlation was above 0.6 (**Fig. 6b**). Three main clusters of NIRS bands were visible and consistent across four environments (e.g., cluster 1: ∼4,000 to 5,000 cm^-1^; cluster 2: ∼5,000 to 7,000 cm^-1^; and cluster 3: ∼7,000 to 10,000 cm^-1^) (**Fig. 6b**). KW had consistently positive correlations with NIRS bands across environments with average correlations of 0.40, 0.36, 0.34, and 0.24 in CS11_WS, CS11_WW, CS12_WS, and CS12_WW respectively; correlation trends decreased as the wavelengths increased (**Fig. 6c**). On average, greater discrepancies in Pearson correlation values between water stressed vs. well-watered environments were observed in CS12, with little difference noticeable in CS11 except in the case of yield (**Fig. 6c**).

**Figure 6.**
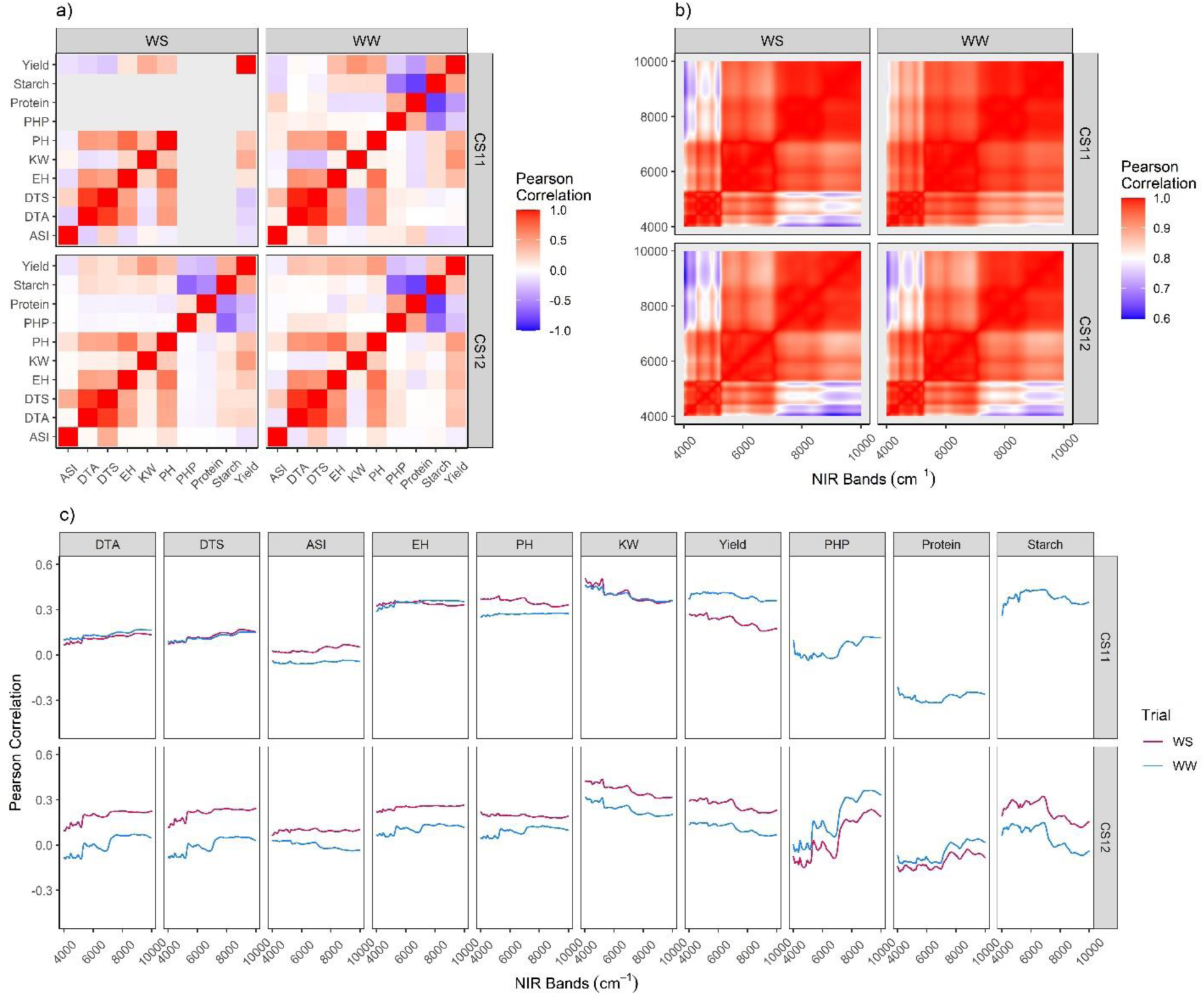
(**a**) The correlations of each of the agronomic traits and kernel composition traits between each other in each environment. (**b**) The correlations of the 3112-band NIR spectrum between each other separated by environment. (**c**) The correlations between each agronomic trait and NIRS bands in each environment.

**Figure 7.**
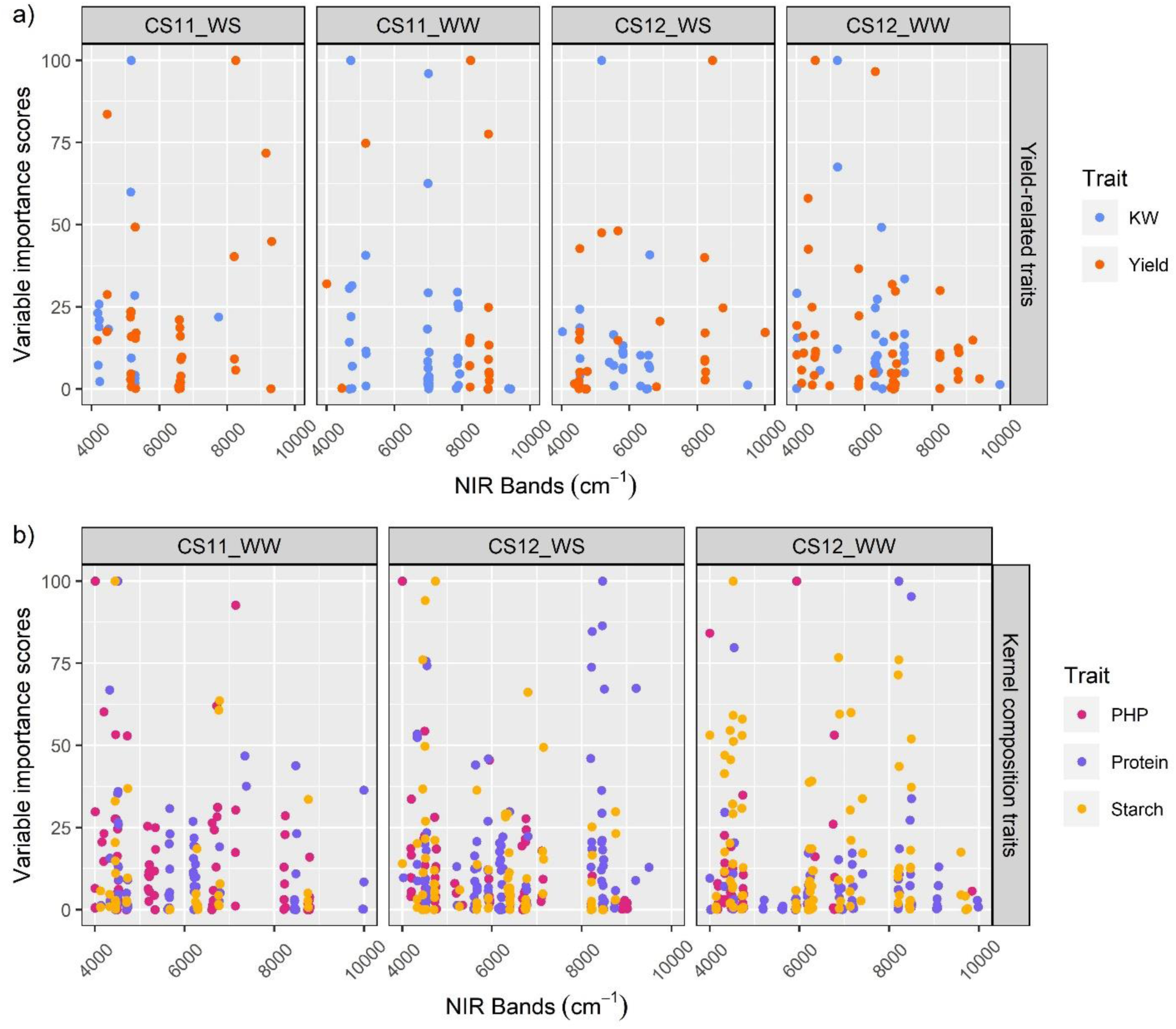
Variable importance scores as calculated by the lasso regression are reported for (**a**) yield-related traits and (**b**) kernel composition traits. Bands with high variable importance are displayed as points with values closer to 100. Data are not reported in CS11_WS due to extreme drought that affected yield in this environment.

### NIRS bands linked to yield-related and kernel composition traits

The lasso algorithm nominated important NIRS bands linked to yield and KW in each environment (**Fig. 7a**). Certain NIRS bands displayed consistency across environments and both traits, such as in the range of ∼5,150 to ∼5,272 cm^-1^ despite the variation in genotypic value distributions of these traits across environments **(Fig. 1b**). R^2^ values ranged from weak (0.072) to strong (0.896) depending on the band for yield-related traits and were generally higher and more consistent for kernel composition traits, ranging from 0.66 to 0.87 (**Supplementary Data 1**). NIRS bands that displayed consistency across environments include ∼4,000 to ∼4,333 cm^-1^ for PHP, ∼4,329 to ∼4,547 cm^-1^ for protein, and ∼6,753 to ∼7,148 cm^-1^ for starch. Variable importance scores and R^2^ values for all bands, traits, and environments are viewable in **Supplementary Data 1**.

### Genome-wide association study

The GWAS nominated 126 unique SNPs (using the Bonferroni-adjusted α value of 0.05) linked to seven agronomic traits and three kernel composition traits across four environments (**Fig. 8**). Of those, 12 SNPs were discovered between two to five times across traits and environments (**Fig. 8; Supplementary Data 1**). Explained phenotypic percent variation by individual loci reached up to 41.2%, 30.7%, 28.2%, and 24.5% for flowering-, height-, yield-related, and kernel composition traits respectively (**Supplementary Data 1**).

**Figure 8.**
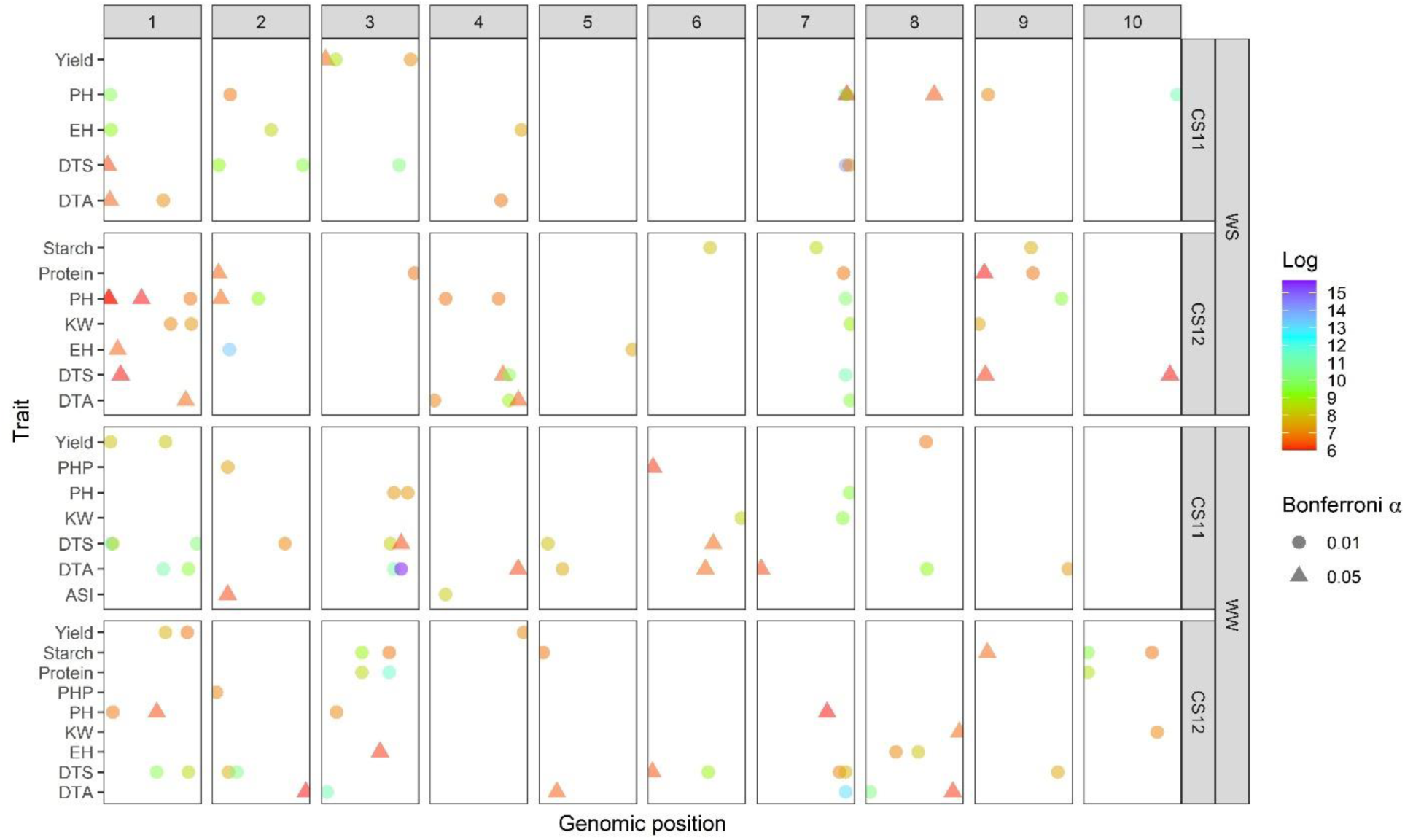
Significant SNPs are displayed separated by chromosome number and environment. Different shapes (in the bottom legend at the right side of the figure) indicate the Bonferroni-adjusted α level at which the SNP was discovered. WW and WS denote well-watered and water stressed environments respectively. The far left side of each block indicates nucleotide position 1 of each of the 10 respective chromosomes.

## Discussion

Improvements in historical crop productivity are largely attributable to traditional plant breeding methods (Evenson and Gollin 2003) and improved agronomic practices (Rizzo et al. 2022). Recent improvements of classical techniques are due in part to the integration of genomic selection into breeding paradigms. Phenomic selection approaches may further improve this paradigm both through increased measurement quantity leading to more accurate breeding value estimations (Lane and Murray 2021), as well as through better understanding and modeling of genetic relationships based on how genotypes interact with the environment. Given the newness of NIRS-based phenomic selection approaches, these have yet to be integrated into breeding programs. The present study provides additional applicability of near infrared spectroscopy measurements (NIRS), an already-prevalent metric for grain quality parameters, to supplement genomic prediction and produce more informed selection decisions across traits using phenomic data. This builds on the body of knowledge shown by other studies on the potential value of NIRS to predict agronomic field performance (Brault et al. 2022; Rincent et al. 2018; Weiß et al. 2022), however the present study also highlights limitations of the technique. Variance component estimation along all 3112 NIRS bands in each of the four environments (CS11_WS, CS11_WW, CS12_WS, CS12_WW) demonstrated that genotypic variance (pedigree) accounted for between 32 – 79% of variation with moderate to high repeatability (**Fig. 2a**). This is consistent across wavelengths and exceeds the explained genetic variance of NIRS reported by Zhu et al. (2021a) and (2022) in soybean and wheat respectively. However, genetic variation and repeatability are dependent on the diversity of germplasm and environments screened, as well as the accuracy of the measurement instrument (NIRS); the present study has no basis for direct comparisons with Zhu et al. (2021a, 2022) or any other study in this regard.

Here we used whole kernel intact samples as opposed to ground homogenized samples, which have much greater throughput and practicality but can lower prediction ability, due to the shallow depth of light penetration into the sample, lack of sample homogenization, and inconsistent light scattering. This may have been a critical limitation in the present study. Spielbauer et al. (2009) note that due to the kernel size of maize being larger than many cereal crops, there is asymmetry in the distribution of kernel composition traits such as protein, starch, and oil in the kernel, which can produce differing NIR spectra on whole kernels when the germinal or abgerminal sides of the kernel are scanned (Heffner et al. 2011; Orman and Schumann 1992; Weinstock et al. 2006). A potentially insightful follow-up to this study might be to scan finely ground kernel samples with NIRS to see if the scanning methodology can “rescue” the predictive ability of the phenomic prediction models (M2) using the same sample genetics and growing environments. Alternatively, to avoid the laborious process of grinding samples, hyperspectral imaging via novel methods developed by Varela et al. (2022) could allow researchers to obtain accurate estimations of kernel composition traits nondestructively, in addition to obtaining morphometric features such as width and length of the kernels. Additional limitations to the present study could include the small sample size of 145 hybrids common across all four growing environments as well as limited effects of G×E owing to similarities between WS and WW environments across years.

Overall patterns of the percent variation explained were similar across the four environments, but the magnitude varied, suggesting the NIR spectra could delineate environmental differences, though the disparities were relatively minor (**Fig. 2b**). Overall, results indicated that NIR spectral bands in this study have potential to serve as a heritable data source to be used in evaluation of agronomically important traits for tested and untested genotypes in observed and unobserved environments.

### Across-Environment Prediction

For flowering-, height– and yield-related traits, genomic prediction (M1) performed better than phenomic prediction alone and similarly to genomic plus phenomic prediction (M3) in unobserved environments (e.g., CV0 and CV00). This indicated that traits measured with relatively high heritability in diverse environments in this study could be sufficiently predicted by genomic data (flowering– and height-related traits), and phenomic data added little value (**Fig. 4**). The latter point contrasts with the findings of Rincent et al. (2018), in which NIR spectra from wheat leaf tissue collected in one environment generated an estimated covariance matrix capable of accurate yield predictions in a weakly correlated separate environment (located ∼500 km apart). Future studies are warranted to determine whether NIRS-based phenomic prediction is suitable as environments become more disparate. Lane et al. (2020) demonstrated the utility of using a functional regression model to predict unobserved hybrids in two out of four environments (predicting 20% of hybrids unobserved in two environments but observed in the remaining two), reporting prediction ability of 0.87 when predicting unknown hybrids in WS while observing these hybrids in WW. The NIRS BLUP model in the same prediction scenario fared worse, reporting a prediction ability of 0.07. Despite many overlapping samples, direct comparisons between the present study and the findings of Lane et al. (2020) are difficult however, as the number of hybrids used for training and testing, structure of the NIRS covariance matrices, and cross-validation methods were different between both studies.

### Within-Environment Prediction

Within-environment prediction using all available hybrids in each environment (199 in CS11_WS, 270 in CS11_WW, 319 in CS12_WS and 220 in CS12_WW respectively) revealed that M1 (genomic only) and M3 (genomic and phenomic) performed similarly for prediction of flowering– and plant height-related traits for within-environment prediction, indicating that the genomic relationship matrix was also more predictive for these scenarios. Overall, genomic and phenomic data together (M3) generally performed similarly or slightly better than genomic data (M1) alone in prediction of yield-related traits for untested genotypes in tested environments (**Fig. 5; Supplementary Data 1**). Thus, within environments, combining near-infrared spectroscopy (NIRS) data with genomic data can enhance the accuracy of predicting grain yield in maize, but not to the extent shown in other studies. Both spectral characteristics of the maize kernels and underlying genetic properties of each hybrid appeared to be viable data sources to predict kernel weight and grain yield in maize for untested genotypes in observed single environments, though the gains in prediction ability afforded by the phenomic data were modest.

### Prediction of kernel composition traits

Among the best PS-predicted results were kernel composition traits (**Supp.** Figs. 3**, 4**). However, we had concerns that because these trait values were obtained from NIRS data and NIRS data were used to develop the predictions, there might be bias in the results. **Supp.** Figs. 2a and 2b indicate the phenomic kernel explained the vast majority of variance for both across– and within-environment phenomic (M2) and combined genomic/phenomic (M3) prediction of all three composition traits. For across-environment prediction of tested (CV2) and untested (CV1) hybrids in known environments, compositional trait prediction abilities of both phenomic (M2) and combined genomic/phenomic (M3) prediction were exceptionally high (all above 0.91). Interestingly, despite the high degree of explained experimental variation by the phenomic data (**Supp.** Fig. 2a), prediction abilities of tested (CV0) and untested (CV00) hybrids in unknown environments were not as robust (**Supp.** Fig. 3). In CV0 and CV00 for kernel composition traits, M3 displayed a notable advantage over genomic (M1) or phenomic (M2) prediction models alone. The distinguishing factor in prediction model performance for these kernel composition traits using NIRS data appeared to be the structure of the training data set, as prediction of unknown hybrids in known single environments (CV1-W) again revealed remarkably high accuracy values (all above 0.91) for all three traits across each of the three environments for which kernel composition values were obtained (**Supp.** Fig. 4). Future studies are warranted to use NIRS-derived phenomic prediction matrices to predict kernel composition traits obtained via wet chemical analysis.

### Genome-wide association study

Several GWAS hits for flowering-related traits were discovered, and of those a GWAS hit (chr8:113842713 bp) which was discovered for DTA in CS12_WW, is around ∼9 kb away from the gene ZCN8. The major role of ZCN8 in maize is to regulate the timing of flowering in response to environmental cues (Meng et al. 2011). The ZCN8 pathway interacts with other genes in the flowering pathway such as ZmMADS69 (Liang et al. 2019). One of the GWAS hits (chr3: 160404839), discovered for DTA and PH in CS11_WW, is ∼1.4 kb away from ZmMADS69. ZmMADS69 directly binds to the promoter region of ZCN8 and activates its expression, suggesting that ZmMADS69 and ZCN8 interact with each other and form a complex that is important for the regulation of flowering time in maize (Liang et al. 2019). This gene was not previously reported by Farfan et al. (2015) in this population.

One GWAS hit (chr9: 17005312 bp) discovered for protein in CS12_WS is ∼5.5 kb away from the *sh1* gene. In maize, this gene encodes the enzyme ADP-glucose pyrophosphorylase (AGPase), which is involved in the initial step of starch biosynthesis in the maize kernel. AGPase is responsible for converting glucose-1-phosphate and ATP to ADP-glucose, which is the primary substrate for starch synthesis. A functional *sh1* gene typically results in the production of a normal amount of AGPase protein, which in turn leads to the normal biosynthesis of starch in the kernel (Chourey and Nelson 1976; Priess 1982). It is important to point out that no sweet corn inbreds (or hybrids) were included in this study. Therefore, it was somewhat surprising that another GWAS hit (chr3: 208042962), which was also discovered for protein in CS12_WS, is ∼8.4 kb away from *sh2* gene. The *sh2* gene is a variant of the wild type *sh1* gene, and it is associated with a higher level of soluble sugars and a lower level of starch in the maize kernel. The *sh2* gene encodes a variant of AGPase that has a higher affinity for ATP than the AGPase encoded by the *sh1* gene, resulting in the preferential synthesis of soluble sugars such as sucrose and fructose rather than starch (Doehlert and Kuo 1990). Another GWAS hit (chr9: 21875116 bp), which was discovered for starch in CS12_WW, is ∼1.4 kb away from *wx1* gene. The *wx1* gene in maize encodes the granule-bound starch synthase (GBSS) that determines the amount and type of starch synthesized in the kernel by means of amylose and amylopectin synthesis (Klösgen et al. 1986). We are unaware of any waxy genotypes in this panel. Another GWAS hit (chr10: 6530030 bp), which was discovered for protein and starch in CS12_WW, is ∼5.2 kb away from *du1* gene. The *du1* gene has a significant effect on the starch content and composition in maize kernels; mutations in this gene result in a dull phenotype with reduced endosperm vitreousness and altered starch properties, leading a reduction in the total starch content and an increase in the proportion of amylose in the starch (Gao et al. 1998). There is some evidence to suggest that the *du1* gene may also influence protein accumulation in the endosperm; the protein content in dull endosperms were lower than that in normal endosperms, possibly due to changes in the balance between starch and protein synthesis in the developing kernel (Cao et al. 1999). Our GWAS hit has negative and positive effect sizes for protein and starch respectively, in accordance with the previously reported results.

Interestingly, five GWAS hits (chr1: 22889335 bp, ch1: 166667449 bp, chr3: 160404839 bp, chr4: 32497838 bp and chr7: 164955163 bp) were discovered for both flowering– and plant height-related traits while three loci were discovered for protein and starch across environments (one of which is the previously mentioned hit at chr10: 6530030 bp). Notably, one locus (chr7: 164955163) was discovered five times (DTS for CS11_WS, CS12_WS, and CS12_WW, and PH for CS11_WS and CS12_WS), and this locus was reported by Farfan et al. (2015) as linked to PH, DTA, and DTS. Our results agree with a previous finding that an interplay between plant height and flowering time by one locus was captured in a stressed environment for maize such as Texas (Adak et al. 2021a; Farfan et al. 2015). Identifying the underlying genetic mechanisms of variation occurring across environments may help to better evaluate the output of prediction models containing both phenomic and genomic data. Thus, success in selection can be performed not only based on black box output of prediction models, but also in the context of plant biology, resulting in competent selection of individuals in breeding populations. However, in total, there was far less heritable genetic variation explained by the GWAS loci detected than by repeatability predictions and genomic selection results.

### Applicability of genomic and phenomic data for plant breeding

As an alternative or supplement to genomic prediction, phenomic prediction uses spectral signals instead of molecular markers. As reflectance spectra are representative of tissue biochemical properties, which are genetically controlled, relationship matrices derived from plant spectra can capture genetic signals (Brault et al. 2022) but also likely environmental and genome×environmental signals. The advent of affordable remote sensing technologies have ushered phenomics into an era of low-cost and higher-throughput data collection, in which spectral data sets (RGB, multispectral, or hyperspectral) enable calculation of vegetation indices that can parameterize predictive models that inform selection decisions (Adak et al. 2021b; Galán et al. 2020; Hernandez et al. 2015; Montesinos-López et al. 2017; Rutkoski et al. 2016). However, these approaches and demonstrations are comparatively new versus more extensively investigated genomic approaches; therefore, new methods and case studies are needed to build bodies of knowledge that improve these methods. Brault et al. (2022) demonstrated that similarities existed between NIRS and genomic relationship matrices across tissues or years in grapevine. These authors, like many others reported to date, found that despite modest genetic variance estimated from NIRS, phenomic prediction was comparable to that of GP.

The present study demonstrated that inclusion of NIRS with genomic relationship matrices has potential to improve prediction abilities of unknown genotypes in both characterized and uncharacterized environments (CV1 and CV00 respectively) for specific traits such as KW in CV00, CV1, and CV2, however genomic prediction (M1) frequently surpassed the predictive ability of phenomic (M2) or combined genomic/phenomic prediction (M3), indicating its continued merit in plant breeding. The cross-validation schemes in this report mimic real-world scenarios faced by plant breeders and demonstrate that more informed models can be generated using readily available spectroscopy platforms. An advantage of phenotypic measurement technologies such as NIRS is the ability to record environment-specific and genetic by environment-specific metrics with the intent of capturing phenotypic plasticity of the same genotypes grown in disparate environments. Given the sensitivity of genomic selection to training/testing set relatedness (Brauner et al. 2020; Olatoye et al. 2020; Riedelsheimer et al. 2012; Weiß et al. 2022; Zhu et al. 2021b), phenomic prediction has shown merit in prediction of diverse breeding material with varying degrees of relatedness (Weiß et al. 2022). Xhu et al. (2021b) and Brauner et al. (2020) reported decreasing trends in genomic prediction ability as the training set composition moved from full-sib to half-sib to unrelated families in *Glycine max* (L.) Merr. and maize, respectively. The diversity of the hybrid panel in the present study may also have played a role in the ability for models to predict across environments, and serves as a key distinction between the elite wheat varieties and association mapping panel of poplar reported by Rincent et al. (2018). Though NIRS-based phenomic prediction alone tended to underperform strictly genomic prediction in this study for flowering– and plant height-related traits in this study, there may still be advantages to phenomic approaches for plant breeders owing to its rapid turnaround time and fixed costs vs. the consumable nature of preparing samples for genotyping. This will be increasingly true where NIRS pipelines are more automated, including NIRS integrated on combine harvesters. As a result, phenomic selection may enable increased selection accuracy with existing population sizes or selection intensity if it allows more lines and hybrids to be screened. It is important to note that using NIRS on kernels and grain before planting would likely be inappropriate in the case of hybrid maize, where the kernel would represent the inbred mother plant and not the genetic potential of the hybrid that will emerge from the seed. In contrast, using the harvested grain, assuming no xenia effect which is reasonable in this study (Bulant et al. 2000), will result in estimations of yield from the plants this grain was selected from. In the case of mechanical (e.g., combine) harvested yield trials, this is unlikely to provide any substantial speed advantages regarding mid-growing season selection decisions, but the enhancement in prediction ability afforded by NIRS data could reduce the amount of screening required in subsequent seasons because genomic/phenomic prediction results can be implemented for either positive or negative selection especially in untested environments.

Prior studies have demonstrated an array of statistical treatments for NIRS in PS since most models applicable to GS can be adapted to PS (Robert et al. 2022b). Functional regression, in which observations are functions outlined on a continuous domain, has also been implemented in PS for yield prediction, leading to greater parsimony among regression models and higher computational efficiency (Lane et al. 2020; Montesinos-López et al. 2018; Montesinos-López et al. 2017; Morris 2015). Variable selection techniques such as lasso and BayesB can reduce multicollinearity and high dimensionality characteristic of NIRS data, where large numbers of wavelengths supersede the number of observations (Lane et al. 2020; Robert et al. 2022b). Variable selection in PS can improve interpretability of findings as certain predictors (in the case of this research, NIRS bands) could show higher heritability than others (Ferragina et al. 2015; Galán et al. 2020). However, Aguate et al. (2017) and Galán et al. (2021) both reported losses in prediction accuracies when less than the full wavelength spectrum (hyperspectral data in these cases) was used. Assessment of variable importance in this study among NIRS bands for phosphorus, protein, and starch (**Fig. 7**) via the lasso regression highlighted wavelengths such as ∼4,000 to 4,333 cm^-1^, 4,329 to 4,547 cm^-1^, and ∼6,753 – 7,148 cm^-1^ that were predictive of these traits respectively. The lasso-selected ranges overlap with previously reported reference ranges for protein and starch in fava bean (Wang et al. 2014). As noted by Zhu et al. (2021a), determining a smaller group of predictive bands creates the potential for production of lower-cost spectrometers that omit non-predictive or repetitive bands.

### Future Directions

The push to explore the predictive nature of agricultural phenomic data has led to the adoption of unoccupied aerial systems (UAS, or drones) to capture temporal, also called longitudinal, phenomic data at scale, which enables the elucidation of genotypic variation across a growing season and potentially well before harvesting. In contrast, genomic information and NIRS of grain represent single measurements (and end of season for NIRS), which is a primary limitation of the present investigation. Future studies are warranted that take advantage of NIRS, RGB, multispectral, and even hyperspectral relationship matrices in attempting to perform prediction of agronomically relevant traits using novel statistical methods to handle large numbers of predictors throughout growth.

### Conclusions

Overall, the use of NIRS data along with genomic data can improve the prediction of agronomic traits of untested genotypes in unobserved environments in maize. Genomic-only prediction was surprisingly more suited to across-environment prediction while combined genomic/phenomic prediction performed best most often for within-environment prediction. Further study of integration of phenomic and genomic data will lead to more accurate and efficient prediction models, reduced need for field-based phenotyping, and an accelerated breeding process. The combination of NIRS and genomic data can overcome some of the limitations of each individual technique, providing a more comprehensive understanding of the genetic and physiological factors that contribute to both agronomically important and research-specific traits in maize such as grain yield, plant height, flowering time, and kernel composition traits, which is fitting especially for breeding programs targeting genetic gains and developing adapted germplasm in their respective governing organizations’ local environments.

## Statements and Declarations

### Funding

Financial support for this research has been provided by: USDA–NIFA–AFRI Award Nos. 2020-68013-32371, and 2021-67013-33915, USDA–NIFA Hatch funds, Texas A&M AgriLife Research, the Texas Corn Producers Board, the Iowa Corn Promotion Board, and the Eugene Butler Endowed Chair in Biotechnology. AJD was supported by the Texas A&M University/Association of Former Students (TAMU/AFS) Graduate Merit Fellowship and the National Science Foundation (NSF) Graduate Research Fellowship (GRFP), and AA was supported by a fellowship from Republic of Turkey, Ministry of National Education and Ministry of Agriculture and Forestry. We would like to acknowledge Colby Bass and Stephen Labar for their agronomic and technical support; graduate and undergraduate students and high school employees of the Texas A&M Quantitative Genetics and Maize Breeding Program for their time and dedication in maintaining fields and collecting phenotypic data. BioRender.com was used in part to create the graphical abstract.

### Competing Interests

The authors declare there are no competing interests.

### Ethics Approval

This research did not use any research materials involving human or animal subjects.

### Author Contributions

All authors contributed to the conceptualization and design of the study. Data curation and analysis were performed by AJD and AA with guidance from DJ. AJD and AA wrote the first draft of the manuscript, and SCM, DJ, NDW, DC, and WR each contributed to revising and editing of previous manuscript versions. All authors approved the final version of the manuscript.

### NSF Statement

Any opinion, findings, and conclusions or recommendations expressed in this material are those of the authors(s) and do not necessarily reflect the views of the National Science Foundation.

## Authors and Affiliations

**Interdisciplinary Graduate Program in Genetics and Genomics (Department of Biochemistry and Biophysics), Texas A&M University, College Station, TX 77843-2128, USA** Aaron DeSalvio (ORCID: 0000-0003-1818-4699)

Department of Soil and Crop Sciences, Texas A&M University, Agronomy Field Lab 110/111, College Station, TX, 77843, USA

Alper Adak (ORCID: 0000-0002-2737-8041), Seth C. Murray (ORCID: 0000-0002-2960-8226), Noah D. Winans (ORCID: 0000-0003-4185-5665), Daniel Crozier (ORCID: 0000-0002-3101-235X), William Rooney (ORCID: 0000-0001-7953-1856)

Department of Agronomy, University of Florida, Gainesville, FL, 32611, USA Diego Jarquín (ORCID: 0000-0002-5098-2060)

## Data Availability

An annotated script of the R code used in this research can be accessed via GitHub (https://github.com/ajdesalvio/Maize-NIRS-GBS.git). Supplementary Data 1 (Supplementary_Data_1.xlsx) contains prediction results, GWAS results, and variable importance scores for NIRS bands. Files necessary to run the R script and reproduce the prediction results are available in the CSVs.zip folder and the SNP60000.hmp.zip folder. Supplementary figures are viewable in the Supplementary_Figures.pdf file.

## Notes

https://github.com/ajdesalvio/Maize-NIRS-GBS.git

### Competing Interest Statement

The authors have declared no competing interest.

